# Naked mole-rat cortical neurons are resistant to acid-induced cell death

**DOI:** 10.1101/260901

**Authors:** Zoé Husson, Ewan St John Smith

**Author notes:** Corresponding author: Ewan St. John Smith, Department of Pharmacology, University of Cambridge, Tennis Court Road, Cambridge, CB2 1PD, United Kingdom. Co-author contact details: Zoé Husson, Department of Pharmacology, University of Cambridge, Tennis Court Road, Cambridge, CB2 1PD, United Kingdom.

## Abstract

Regulation of brain pH is a critical homeostatic process and changes in brain pH modulate various ion channels and receptors and thus neuronal excitability. Tissue acidosis, resulting from hypoxia or hypercapnia, can activate various proteins and ion channels, among which acid-sensing ion channels (ASICs) a family of primarily Na^+^ permeable ion channels, which alongside classical excitotoxicity causes neuronal death. Naked mole-rats (NMRs, *Heterocephalus glaber*) are long-lived, fossorial, eusocial rodents that display remarkable behavioral/cellular hypoxia and hypercapnia resistance. In the central nervous system, ASIC subunit expression is similar between mouse and NMR with the exception of much lower expression of ASIC4 throughout the NMR brain. However, ASIC function and neuronal sensitivity to sustained acidosis has not been examined in the NMR brain. Here, we show with whole-cell patch-clamp electrophysiology of cultured NMR and mouse cortical and hippocampal neurons that NMR neurons have smaller voltage-gated Na^+^ channel currents and more hyperpolarized resting membrane potentials. We further demonstrate that acid-mediated currents in NMR neurons are of smaller magnitude than in mouse, and that all currents in both species are fully blocked by the ASIC antagonist benzamil. We further demonstrate that NMR neurons show greater resistance to acid-induced cell death than mouse neurons. In summary, NMR neurons show significant cellular resistance to acidotoxicity compared to mouse neurons, contributing factors likely to be smaller ASIC-mediated currents and reduced NaV activity.

## Background

Acid-sensing channels (ASICs) are ion channels of the ENaC/Deg superfamily and most subunits are activated by extracellular protons [1,2]. Six different ASIC subunits are encoded by 4 ASIC genes (ASIC1a, ASIC1b, ASIC2a, ASIC2b, ASIC3 and ASIC4), which assemble as homo- or heterotrimers [3]; neither ASIC2b nor ASIC4 form proton-sensitive homotrimers. ASICs are primarily permeable to Na^+^, although ASIC1a homomeric channels are also Ca^2+^ permeable [2]. In the central nervous system (CNS), neurons have been shown to primarily express ASIC1a homomers and heteromers of ASIC1a/2a and ASIC1a/2b [4–8], where they have been demonstrated to have key roles in synaptic plasticity [9–11] and fear conditioning [12–14], as well as being major players in neuronal death resulting from brain ischemia [5,15–17], and neurodegenerative diseases [18–20].

Regulation of brain pH is a highly complex and important process [21]. Brain tissue acidosis can result either from an increase in tissue partial pressure of carbon dioxide (PCO2) during hypercapnia, or from the accumulation of the byproducts of anaerobic metabolism, such as lactate and protons, during hypoxia [22]. During periods of tissue acidosis, activation of ASICs by extracellular acidification is worsened by the release of allosteric modulators such as lactate [23], spermine [16] and arachidonic acid [24,25]. In addition to the activation of the Ca^2+^ permeable ASIC1a channel [15,16], a drop in pH also modulates the activity of numerous others ion channels, including voltage-gated ion channels [26–29] and glutamate receptors [30,31], therefore leading to disturbance in ion homeostasis, excitotoxicity and ultimately neuronal death [5,15,16].

Naked mole-rats (NMRs, *Heterocephalus glaber*) are subterranean rodents belonging to the Bathyergidae African mole-rat family found in East Africa [32]. Unusually for a mammal, NMRs are eusocial [33,34]. However, NMRs also display a range of remarkable physiological peculiarities, which is beginning to make a significant impact on biomedical research [35]. The unusual physiology of the NMR includes: extreme longevity with no increased risk of death with ageing [36], an apparent absence of age-related neurodegenerative disorders [37,38], resistance to cancer [39–41], insensitivity to certain noxious and irritant stimuli [42–45] and hypoxia/hypercapnia resistance resulting from altered NMDA receptor function and an ability to utilize fructose as an energy source [46–49]. It is striking that NMRs are resistant to many pathological conditions known to involve ASICs. Recordings of ASIC-mediated currents in dorsal root ganglion (DRG) sensory neurons demonstrated an increased frequency and magnitude of ASIC responses in NMR neurons compared to mouse neurons [43], with APETx2, an inhibitor of ASIC3-containing ASICs, demonstrating a key role for ASIC3, even though nmrASIC3 does not appear to form functional homotrimers [50]. Previously we mapped out ASIC expression in different NMR brain regions and observed similar expression between mouse and NMR, a key exception being much lower ASIC4 levels throughout the NMR brain [51], however, no one has yet studied the function of ASICs in NMR brain neurons.

In this study, we investigated acid-induced currents in mouse and NMR neurons using whole-cell patch clamp recording of cultured neonatal hippocampal and cortical neurons. We find that NMR neurons have ASIC-mediated currents of significantly smaller peak current amplitude than those recorded from mouse neurons and that NMR neurons are resistant to acid-induced cell death. Overall, these results suggest that the reduced acid-induced cell death in NMR neurons may be neuroprotective.

## Methods

### Animals

All experiments were conducted in accordance with the United Kingdom Animal (Scientific Procedures) Act 1986 Amendment Regulations 2012 under a Project License (70/7705) granted to E. St. J. S. by the Home Office; the University of Cambridge Animal Welfare Ethical Review Body also approved procedures. Breeding couples of 1 male and 2 female C57/bl6 mice were conventionally housed with nesting material and a red plastic shelter; the holding room was temperature-controlled (21 °C) and mice were on a normal 12-hour light/dark cycle with food and water available *ad libitum*. Naked mole-rats were bred in house and maintained in a custom-made caging system with conventional mouse/rat cages connected by different lengths of tunnel. Bedding and nesting material were provided along with running wheels and chew blocks. The room was warmed to 28 °C and humidified, with a heat cable to provide extra warmth running under 2–3 cages, and red lighting (08:00–16:00) was used.

### Neuronal cultures

P0-P2 mice and P0-P5 naked mole-rats were used to prepare cortical and hippocampal neuronal cultures. Multiple pups (2-4) were used to prepare a single culture. Following decapitation, heads were immediately placed in dishes containing ice-cold Hank’s Balanced Salt Solution (HBSS) solution (20 mM HEPES, 30 mM glucose in HBSS, Life Technologies). Brains were removed, transferred to a new dish and the two hippocampi and cortices were isolated. Tissues were subsequently incubated in an enzymatic digestion solution: 2 mg/mL papain (Worthington Biochemical Corporation) in Hibernate-Ca^2+^ solution (Brain Bits), activated by 0.5 mM Glutamax (Life Technologies) at 37 °C for 30 min in a 5 % CO2 incubator. The digestion solution was then replaced by HBSS solution supplemented with DNAse I (250 Kunitz units/mL, Sigma Aldrich) and tissues were slowly triturated (5-7 times) using a P1000 pipette. Neuronal suspensions were filtered through a 100 µm nylon cell strainer (Corning) to remove non-dissociated pieces of tissues before centrifugation for 5 mins at 1100 rpm at room temperature. Supernatants were discarded and the pellets carefully resuspended in HBSS solution. After further centrifugation for 5 min 1100 rpm at room temperature, pellets were resuspended in MEM/HS solution: 10 % heat-inactivated horse serum (Life Technologies), 2 mM glucose, 0.0025 % Glutamax, and 0.2 mg/mL primocin (InVivogen). Hippocampal and cortical neurons were plated on 35 mm plastic dishes (Fisher Scientific), previously coated with 100 mg/mL poly-L-lysine (Sigma-Aldrich), rinsed with water and dried, at a density of 300 000 cells/mL (2 mL/dish). After a 4-hour incubation in a 37 °C / 5 % CO2 incubator, the MEM/HS solution was removed and the dishes were flooded with Neurobasal/B27 solution (1X B27 Supplement, 0.0025 % Glutamax, and 0.2 mg/mL primocin). Naked mole-rat neurons were kept at 33 °C in 5 % CO2 incubator, whereas mouse neurons were kept at 37 °C in a 5 % CO2 incubator; this is due to NMRs being cold-blooded and NMR cells do not withstand 37 °C for long periods of time [52]. Half of the medium was exchanged for fresh medium every 2-3 days until the cultures were used for experiments.

### Electrophysiology

Hippocampal and cortical neurons from mouse and NMR were used for whole-cell patch-clamp recordings at 9-12 days *in* vitro (DIV9-12). Recordings were performed at room temperature using the following solutions: extracellular (in mM) – 140 NaCl, 4 KCl, 2 CaCl2, 1 MgCl2, 4 glucose, 10 HEPES, adjusted to pH 7.4 with NaOH and 300–310 mOsm with sucrose; intracellular (in mM) – 110 KCl, 10 NaCl, 1 MgCl2, 1 EGTA, 10 HEPES, 2 Na2ATP, 0.5 Na2GTP, adjusted to pH 7.3 with KOH and to 310–315 mOsm with sucrose. Acidic extracellular solutions were made using MES (pH 5.0). Patch pipettes were pulled (Model P-97, Flaming/Brown puller; Sutter Instruments,) from borosilicate glass capillaries (Hilgenberg GmbH) and had a resistance of 6-10 MΩ. Data were acquired using an EPC10 amplifier and Patchmaster software (HEKA). Whole-cell currents were recorded at 20 kHz, pipette and membrane capacitance were compensated using Patchmaster macros, and series resistance was compensated by >60 %. Cell capacitances and resting membrane potentials were measured just after cell opening in whole-cell configuration. To study macroscopic voltage-gated currents, a standard voltage-step protocol was used whereby cells were held at −120 mV for 200 msecs before stepping to the test potential (−80 mV - +65 mV in 5 mV increments) for 50 msecs, returning to the holding potential (−60 mV) for 200 msecs between sweeps. In some experiments, tetrodotoxin (300 nM, Alomone Labs) was perfused for 30 seconds before repeating the voltage-step protocol. To measure neuronal acid-sensitivity, cells were exposed to the following protocol: 5 seconds of pH 7.4; 5-seconds of pH 5; and 5 seconds of pH 7.4. ASIC antagonists (100 µM Benzamil, Sigma) were perfused during 30 seconds before applying another 5 second pulse of pH 5. After 90 second wash time with pH 7.4 solution, a 5 second pulse of pH 5 was applied to check for reversal of any block observed. Current amplitude was measured in Fitmaster (HEKA) by taking the maximum peak response and subtracting the mean baseline amplitude in the preceding 50 msec (voltage-gated currents) or ~2.5 sec (ASIC currents); current amplitude was normalized for cell size by dividing by cell capacitance. Using Igor Pro, for each individual cell that underwent the voltage-step protocol, the following equation was fitted to the normalized inward currents:

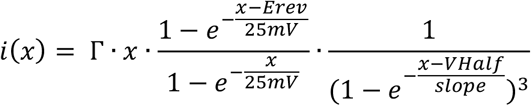

where Erev is the reversal potential; Vhalf the half-activating potential; Γ a constant and x the command potential. Similarly, a Boltzmann equation was fitted to the normalized outward currents:

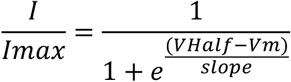

where Vm is the membrane voltage and VHalf the voltage at half-maximal activation. To determine the inactivation time of the ASIC-mediated currents, a single exponential was fitted. Data are expressed as mean ± standard error of the mean (SEM). Using Prism (GraphPad), paired t-tests were used to compare the effects of ASIC antagonists on proton-gated currents within both mouse and NMR neuron datasets; unpaired t-tests were used to compare parameters, such as neuronal resting membrane potential and capacitance and ASIC-mediated current amplitude, between mouse and NMR neuron datasets.

### Acid-induced cell death assays

Mouse and NMR neurons were used to measure acid-induced cell death at DIV9-12. The pH 7.4 and pH 5 extracellular solutions used were the same as those described above for electrophysiology experiments. Neuronal cultures from both mouse and NMR were rinsed twice with 37 °C solution (pH 7.4 or pH 5) and then incubated with pH 7.4 or pH 5 solution for 2 hours at 37 °C. Cultures from both conditions were then rinsed with warm pH 7.4 solution and incubated during 30 min with pH 7.4 solution containing 1.5 mM propidium iodide (PI, Sigma Aldrich) to stain necrotic cells and Hoechst 33342 (dilution 1/2500, Sigma) to label all nuclei. Labelled cultures were imaged using an epifluorescence microscope (Olympus) equipped with a 20X objective (Olympus) and a QImaging camera. To determine the percentage of dead necrotic PI-positive cells, we used the software Fiji to count the total number of cells per field of view by counting nuclei on the Hoechst images, and subsequently counting the number of PI-positive nuclei on the PI images. One to three dishes per condition were used, and three different images per dish were taken. Data were collected from three different cultures and each culture was prepared from multiple animals. A one-way ANOVA test (Prism, GraphPad) corrected for multiple comparisons (Tukey test) was used to compare the percentage of cell death in each field of view at pH 7.4 and pH 5 in both species. Data are expressed as mean ± standard error of the mean (SEM).

## Results

### Basic electrophysiological properties and voltage-gated Na^+^ channel activity differ between NMR and mouse neurons

Electrophysiological recordings from NMR neurons have been performed in both DRG sensory neurons [43,50] and CNS neurons [46,47,53]. However, neuronal activity from NMR CNS has only been recorded in brain slices, in the form of field excitatory postsynaptic potentials, and the basic electrophysiological properties of NMR neurons in hippocampal and cortical cultures have not yet been described.

We first compared the capacitance and resting membrane potential of NMR and mouse neurons from both cortical and hippocampal neuronal cultures (Fig. 1). The capacitance of NMR neurons was significantly smaller than in mouse neurons in both cortical and hippocampal cultures (cortex: 17.27 ± 1.02 pF versus 28.72 ± 2.32 pF for NMR (n = 30) and mouse (n = 24) neurons, respectively; hippocampus: 17.41 ± 1.03 pF versus 38.13 ± 3.27 pF for NMR (n = 18) and mouse (n = 26) neurons, respectively; unpaired two-sided t-tests, **** p < 0.0001, Fig. 1a). Resting membrane potentials were measured as soon as the whole-cell configuration was established and NMR neurons were significantly more hyperpolarized than mouse neurons, in both cortical and hippocampal cultures (cortex: −57.03 ± 2.64 mV versus −44.05 ± 2.84 mV for NMR (n = 30) and mouse (n = 21) neurons, respectively; hippocampus: −55.11 ± 4.79 mV versus - 43.85 ± 2.59 mV for NMR (n = 18) and mouse (n = 26) neurons, respectively; unpaired two-sided t-tests; ** p < 0.01; * p < 0.05, Fig. 1b).

**Fig. 1.**
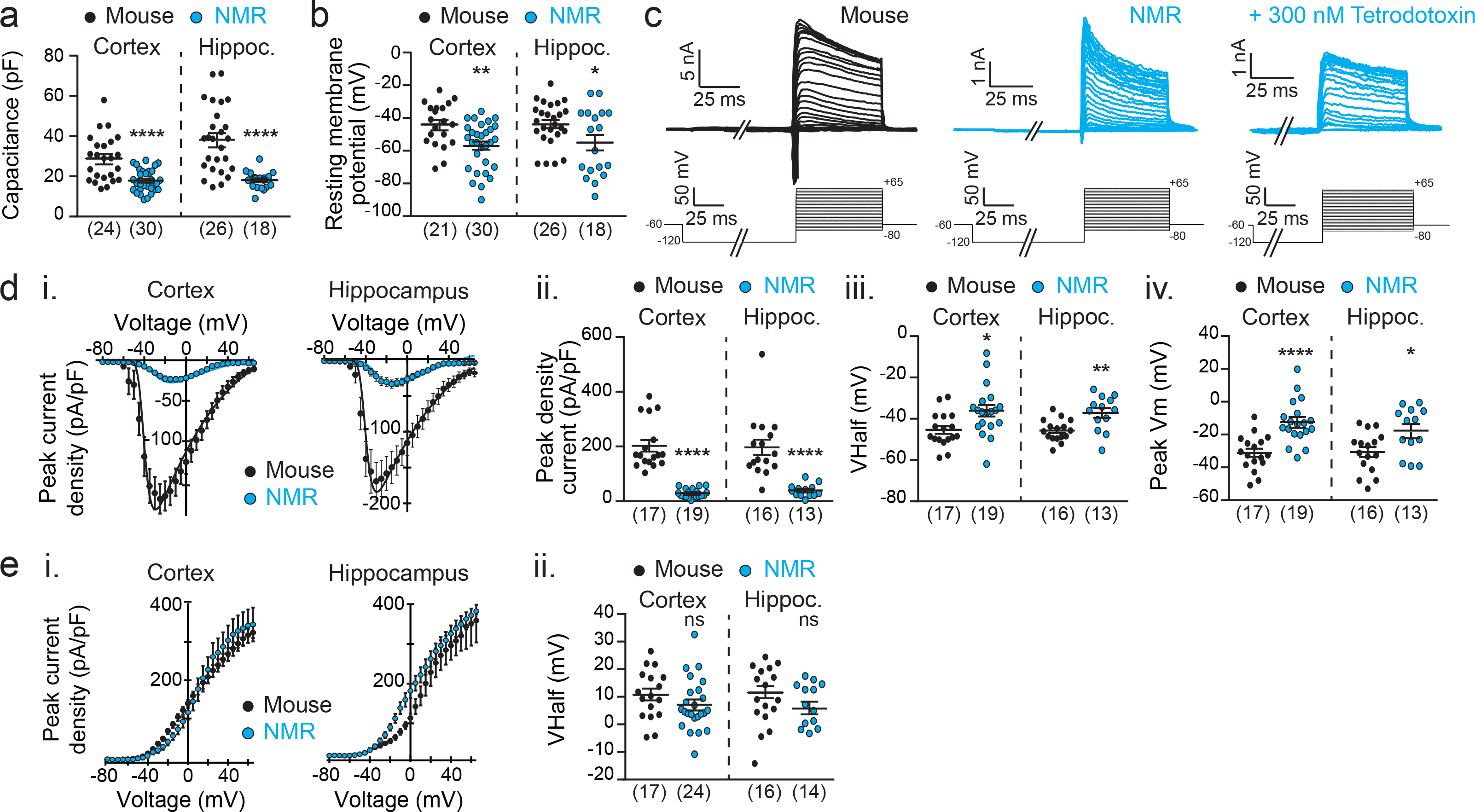
Electrophysiological properties of NMR neurons differ from mouse neurons. NMR cortical and hippocampal neurons have significantly smaller capacitance (**a**) and more hyperpolarized resting membrane potential (**b**) than mouse neurons. **c.** Example of whole-cell patch-clamp recordings made from a mouse (left, black) and a NMR (middle, blue) hippocampal neuron in response to the voltage step protocol and the inhibition of response in NMR neurons by 300 nM TTX (right, blue). Cortical and hippocampal NMR neurons have significantly smaller voltage-gated inward currents (**a, b**), a more depolarized Vhalf (**c**) and a more depolarized peak activation membrane potential Vm (**d**) compared to mouse neurons. **e.** Voltage-gated outward currents in NMR cortical and hippocampal show no significant difference to those in mouse neurons. * p < 0.5; ** p < 0.01; **** p< 0.0001, unpaired t-tests within each structure. Numbers in brackets indicate the number of recorded cells.

We then investigated macroscopic voltage-gated currents in NMR and mouse neurons, using a voltage-step whereby cells were held at −120 mV for 200 msecs before stepping to the test potential (−80 mV to +65 mV in 5 mV increments) for 50 msecs, and returning to the holding potential (−60 mV) for 200 msecs between sweeps (Fig. 1c). Both NMR and mouse neurons showed inward and outward currents (Fig. 1c-e). In some experiments, the voltage-step protocol was run twice, the second time after 300 nM tetrodotoxin (TTX) had been applied for 30 seconds to investigate the contribution of TTX-sensitive voltage-gated Na^+^ channels (NaVs) to the macroscopic voltage-gated inward currents recorded in NMR neurons (Fig. 1c, right panel). In both cortical and hippocampal NMR neurons, the voltage-gated inward currents were fully blocked by 300 nM TTX (n = 8 and n = 2 for cortical and hippocampal neurons, respectively). The fact that no inward current remained after application of 300 nM TTX indicates that only NaVs were activated with our voltage-step protocol and that there was no measurable contribution of voltage-gated Ca^2+^ channels to the inward currents recorded. Moreover, these results indicate that cortical and hippocampal NMR neurons only express TTX-sensitive NaVs.

Strikingly, NMR neurons had a significantly smaller inward peak current density in both cortical and hippocampal cultures (cortex: 28.79 ± 3.83 pA/pF versus 202.40 ± 21.35 pA/pF for NMR (n = 19) and mouse (n = 17) neurons, respectively; hippocampus: 39.00 ± 6.39 pA/pF versus 196.70 ± 28.28 pA/pF for NMR (n = 13) and mouse (n = 16) neurons, respectively; unpaired two-sided t-tests, **** p < 0.0001, Fig. 1d.i-ii). Additionally, the voltage of half-activation (Vhalf) and the peak inward current amplitude potential (peak Vm) were more depolarized in NMR neurons compared to mouse neurons (cortex: Vhalf: −36.12 ± 2.80 mV versus −45.42 ± 1.99 mV and peak Vm: −12.43 ± 3.02 mV versus −31.27 ± 2.61 mV for NMR (n = 19) and mouse (n = 17) neurons, respectively; hippocampus: Vhalf: −37.10 ± 2.34 mV versus −45.71 ± 1.30 mV and peak Vm: - 17.62 ± 4.00 mV versus −30.68 ± 3.09 mV for NMR (n = 13) and mouse (n = 16) neurons, respectively; unpaired two-sided t-tests, **** p < 0.0001, ** p < 0.01; * p < 0.05, Fig. 1d.iii-iv). These results suggest that NMR neurons may be less excitable compared to mouse neurons, with more hyperpolarized resting membrane potentials and smaller voltage-gated inward currents that are activated at more depolarized potentials, i.e. a greater depolarizing stimulus is required to activate NMR voltage-gated inward currents that produce much smaller currents.

By contrast, current-voltage curves for voltage-gated outward currents were similar between NMR and mouse neurons from both cortical and hippocampal neurons (Fig. 1e.i) and Vhalfs were not significantly different (cortex: 7.13 ± 1.91 mV versus 10.77 ± 2.18 mV for NMR (n = 24) and mouse (n = 17) neurons, respectively; hippocampus: 5.15 ± 2.50 mV versus 11.59 ± 2.26 mV for NMR (n = 14) and mouse (n = 16) neurons, respectively; unpaired two-sided t-tests, p = 0.221 and p = 0.094 for cortex and hippocampus respectively, Fig. 1e.ii).

### Acid-induced currents are mediated by ASICs in both NMR and mouse neurons

The expression profile of ASIC subunits is similar in mouse and NMR brains, with the exception of lower levels of ASIC4 throughout the NMR brain [51], results suggesting that functional ASIC-mediated currents should be present in NMR as others have shown in mouse [4,5,7,8,15].

A 5 second pulse of pH 5 was applied to NMR and mouse neurons from both hippocampal and cortical neurons and rapidly activating and inactivating acid-induced responses were recorded in every cell of both species (Fig. 2a). However, the peak current density of acid-mediated responses recorded in NMR neurons was significantly smaller than in mouse neurons, in both hippocampal and cortical neurons (cortex: 15.41 ± 1.82 pA/pF versus 85.66 ± 10.90 pA/pF for NMR (n = 31) and mouse (n = 24) neurons, respectively; hippocampus: 20.54 ± 2.86 pA/pF versus 100.90 ± 17.68 pA/pF for NMR (n = 22) and mouse (n = 26) neurons, respectively; unpaired two-sided t-tests, **** p < 0.0001, *** p < 0.001, Fig. 2b). By contrast, the inactivation time constant of the acid-mediated currents was similar between NMR and mouse neurons (cortex: 0.35 ± 0.012 s versus 0.44 ± 0.056 s for NMR (n = 27) and mouse (n = 18) neurons, respectively, unpaired two-sided t-test, p=0.0516; hippocampus: 0.47 ± 0.06 s versus 0.35 ± 0.04 s for NMR (n = 13) and mouse (n = 19) neurons, respectively; unpaired two-sided t-test, p = 0.0861, Fig. 2c).

**Fig. 2.**
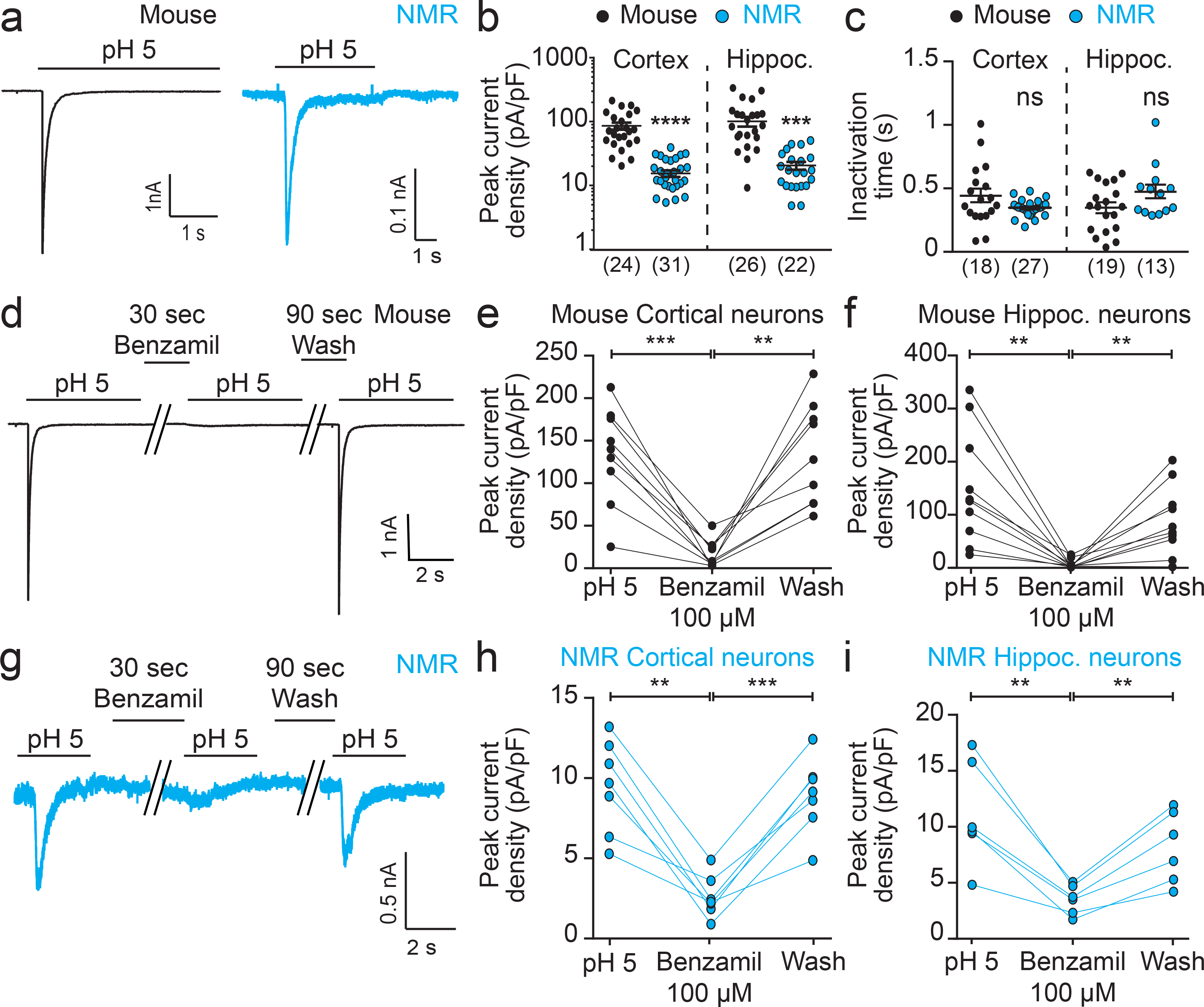
ASICs mediate acid-induced currents in NMR and mouse CNS neurons. **a.** Both mouse (black trace, left) and NMR (blue trace, right) neurons respond to a pH 5 solution with a transient inward current. **b.** Acid-induced currents are of significantly smaller amplitude in NMR neurons compared to mouse neurons. **c.** Inactivation time constants of the acid-induced responses are similar between NMR and mouse neurons. **d.** Example trace of an acid-induced current elicited by a pH 5 solution and the effect of 100 µM benzamil in a mouse cortical neuron. **e-f.** Acid-induced currents in both cortical and hippocampal mouse neurons were reversibly blocked by 100 µM benzamil (n = 9 and n = 10 cortical and hippocampal neurons, respectively). **g.** Example trace of an acid-induced current evoked by a pH 5 solution in a cortical NMR neuron showing inhibition by 100 µM benzamil. **h-i.** In both cortical (n = 7) and hippocampal (n = 6) NMR neurons, acid-induced currents were reversibly blocked by 100 µM benzamil. ** p < 0.01; **** p< 0.0001, one-way ANOVA paired tests, Tukey’s multiple comparison tests. Numbers into brackets indicates the number of recorded cells.

The transient nature of the acid-mediated inward currents in both NMR and mouse neurons is characteristic of ASIC-mediated currents [3] and to confirm the involvement of ASICs we utilized the non-selective ASIC antagonist benzamil [54]. After a first application of pH 5 for 5 seconds, we applied 100 µM benzamil for 30 seconds before a second pH 5 pulse, then followed by a wash period of 90 seconds (Fig. 2d, 2g). In cortical and hippocampal mouse neurons, the acid-induced currents were reversibly blocked by 100 µM benzamil (cortex: pH 5: 133.60 ± 19.01 pA/pF; benzamil: 16.77 ± 5.28 pA/pF; wash: 133.80 ± 19.88 pA/pF (n = 9); hippocampus: pH 5: 151.00 ± 33.64 pA/pF; benzamil: 7.84 ± 2.67 pA/pF; wash: 89.26 ± 20.48 pA/pF (n = 10); one-way paired ANOVA test, Tukey’s multiple comparison test, *** p < 0.001, ** p < 0.01, Fig. 2e-f). This result concurs with previous studies indicating that acid-evoked currents in mouse hippocampal and cortical neurons are mediated by ASICs [4,8,15].

Similarly, in cortical and hippocampal NMR neurons, acid-induced currents were reversibly inhibited by 100 µM benzamil (cortex: pH 5: 9.66 ± 1.09 pA/pF; benzamil: 2.78 ± 0.49 pA/pF; wash: 9.13 ± 0.89 pA/pF (n = 7); hippocampus: pH 5: 11.13 ± 1.88 pA/pF; benzamil: 3.50 ± 0.53 pA/pF; wash: 8.15 ± 1.30 pA/pF (n = 6); one-way paired ANOVA test, Tukey’s multiple comparison test, *** p < 0.001, ** p < 0.01, Fig. 2h-i). This is the first demonstration of functional ASIC-mediated currents in CNS NMR neurons.

### NMR neurons are resistant to acid-induced cell death

ASICs are involved in acid-induced cell death, so-called acidotoxicity, which can occur during periods of ischemia [5,15,16]. Because NMR neurons exhibit significantly smaller ASIC currents (Fig. 2), we hypothesized that this could be neuroprotective when neurons are in an acidic environment. We exposed neuronal cultures from mouse and NMR cortices to a pH 7.4 or a pH 5 solution for 2 hours at 37 °C. Nuclei were stained by Hoechst 33342 and necrotic (dead) neurons were labeled using propidium iodide (PI) (Fig. 3). Percentages of dead neurons were calculated by counting the number of necrotic PI-positive cells over the total number of Hoechst 33342-positive neurons in the field of view, and a one-way ANOVA test corrected for multiple comparisons (see *Methods*) was used to compare neuronal death at pH 5 between both species (Fig. 3c). At pH 7.4, percentages of dead cells were comparable between mouse and NMR cortical cultures (mouse - pH 7.4: 20.31 ± 3.50 %, n = 18 fields of view; NMR – pH 7.4: 13.31 ± 1.53 %, n = 27 fields of view; from 3 independent experiments), suggesting no differences in neuronal death under basal conditions between NMR and mouse cultures. When incubated for 2 hours in pH 5 solution, mouse cortical neurons exhibited a significantly increased percentage of cell death compared to all other conditions (mouse – pH 5: 60.55 ± 4.87 %, n = 18 fields of view; NMR – pH 5: 22.29 ± 2.30 %, n = 28 fields of view, from 3 independent experiments, p < 0.0001, ****). By contrast, when NMR cortical neurons were exposed to pH 5 for 2 hours, no difference in the level of cell death was observed compared to incubation at pH 7.4. Similar results were obtained using mouse and NMR hippocampal neurons (data not shown). This indicates that NMR neurons are resistant to acid-induced cell death, possibly due to their reduced ASIC currents compared to mouse neurons.

**Fig. 3.**
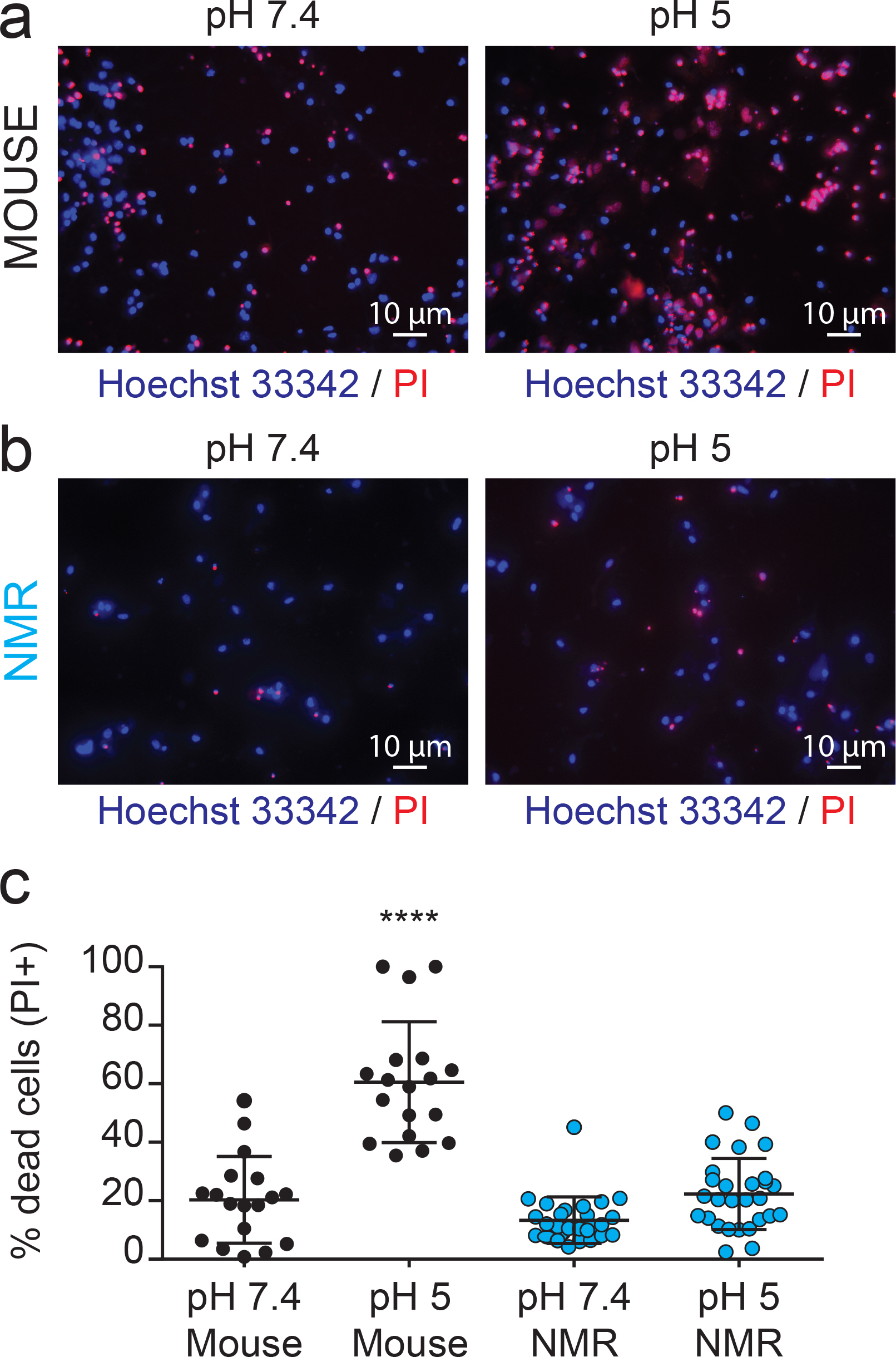
NMR cortical neurons show resistance to acid-induced cell death. Mouse (**a**) and NMR (**b**) cortical neurons were incubated for 2 hours with either a pH 7.4 or pH 5 solution and then stained with Hoechst 33342 to label nuclei and PI to label necrotic dead cells. **c.** Percentages of PI-positive neurons (i.e. dead) at pH 7.4 were similar between mouse and NMR neurons. The percentage of mouse dead neurons was higher when cells were incubated in a pH 5 solution, compared to pH 7.4 condition and compared to the percentage of cell death obtained with NMR neurons at pH 5. NMR cortical neurons did not exhibit any significant difference with regard to cell death when exposed to pH 5.0 compared to pH 7.4. **** p< 0.0001, one-way ANOVA test, Tukey’s multiple comparison test. Percentages of each field of view were used to compare acidotoxicity between species. The number of fields of view analyzed for each condition are the following: n = 18 for mouse neurons (for both pH 7.4 and pH 5), n = 27 and 28 for NMR neurons at pH 7.4 and at pH 5, respectively; obtained from 3 independent cultures for both species.

## Discussion

In this study, we recorded for the first time from cultured NMR brain neurons and described their basic electrophysiological properties (Fig. 1). We showed that NMR neuronal capacitance was smaller than in mouse neurons and that the resting membrane potential of NMR neurons is more hyperpolarized than in mouse neurons. One point to consider is that neurons were recorded from blindly and so we can only broadly compare hippocampal and cortical neurons between species and cannot comment on if these differences occur in all types of neuron or specific neuronal subpopulations, such as interneurons. Although capacitance and resting membrane potential values obtained from mouse neurons were similar to those reported by others using cultured rodent neurons [55,56], for NMR neurons however, previous data from others found their resting membrane potential to be more hyperpolarized (NMR cortical neurons: −57.03 ± 2.64 mV, NMR hippocampal neurons: −55.11 ± 4.79 mV, Fig. 1; NMR hippocampal pyramidal neurons (4 months old): −70.3 ± 6.1 mV, NMR hippocampal dentate granule cells (4 months old): - 75.1 ± 3.6 mV, [53]). Explanations for differences observed in resting membrane potentials between studies are the recording conditions (*in vitro* vs. *in vivo*), the developmental stage of the animal from which neurons are obtained and the constituents of the intracellular and extracellular solutions. When comparing our study with that of Penz and colleagues, who reported more hyperpolarized resting membrane potentials, two factors that could contribute to the difference observed is that we made recordings from cultured neurons, whereas they used brain slices, and secondly, we isolated neurons from neonatal animals, whereas slice recordings were made from animals aged at least 4-months.

A standard voltage-step protocol that has been already successfully used in NMR sensory neurons to predominantly isolate NaV activity [43] was used to measure macroscopic voltage-gated current activity in mouse and NMR neurons. NaV currents recorded from mouse neurons were not different to what have been recorded in similar experimental conditions (for example, mouse cortical neurons at DIV 10-12: Vhalf: −41.62 ± 1.46 mV in [57] versus this study : −45.42 ± 1.99 mV). Similarly, inward currents recorded in NMR brain neurons were also very similar to currents recorded in NMR sensory neurons (NMR cortex: Vhalf: −36.12 ± 2.80 mV; NMR hippocampus: Vhalf: −37.10 ± 2.34 mV; NMR DRG neurons: Vhalf: from −34.2 ± 0.1 mV to −44.6 ± 0.5 mV [43]). However, NMR inward currents significantly differed from mouse currents: the peak current density was significantly smaller and the Vhalf and peak Vm values were more depolarized. These differences in NMR NaV activity likely result from differences in amino acid sequence of NaV subunits and/or differential expression of accessory subunits and warrant further investigation. We also found that addition of 300 nM TTX completely abolished all voltage-gated inward currents demonstrating an absence of TTX resistant NaV subunits in the NMR brain, this result aligns with the observation that the TTX-resistant NaV subunits, NaV1.5, NaV1.8 and NaV1.9 are not expressed in the central nervous system [58]. It should also be noted that although NMR neurons were cultured at 32 °C under hypoxic (3 % O2) conditions, recordings under normoxic, standard laboratory conditions, whereas NMRs live in a hypoxic and hypercapnic environment [59,60], which may influence channel activity *in vivo*.

ASIC activity in brain neurons is now accepted as a key factor in numerous physiological and pathological conditions [61]. However, nothing is known about acid-induced responses in NMR brain neurons, which is of considerable interest considering the behavioral hypoxia/hypercapnia resistance and lack of acid-induced nocifensive behavior display by NMR that are likely adaptations to adapting to a safe, but relatively hypoxic and hypercapnic habitat [35]. Recently, we described ASIC subunit expression in the NMR CNS [62], which is similar to that in the mouse CNS, with the exception of lowered ASIC4 levels in the NMR brain. However, evidence for functional ASIC activity is lacking and is of particular interest in light of our recent finding that nmrASIC3 forms non-functional homomers [50]. Here, we find that NMR brain neurons produce ASIC-mediated currents in response to acid stimulation, in both hippocampus and cortex, as demonstrated by acid-induced responses being fully, reversibly blocked by 100 µM benzamil (Fig. 2). However, the peak current density of NMR ASIC-mediated responses was significantly reduced compared to responses recorded in mouse neurons. The reasons for such reduction in ASIC currents in NMR neurons is not known and additional research is needed to determine what underpins this different, e.g. are regulators of ASIC plasma membrane trafficking different in NMR? Is there a different developmental expression profile of ASICs between mouse and NMR? With regard to the ASIC currents themselves, they have similar decay kinetics in both NMR and mouse neurons (Fig. 2c), which suggests that a similar mixture of ASIC subunits are expressed, as our previous mRNA based analysis suggested [62].

Incubation with a pH 5.0 solution showed that unlike mouse neurons, NMR neurons do not undergo any significant acid-induced cell death (Fig. 3). This is the first demonstration of resistance to acid-induced neuronal death in NMR neurons. In rodent neurons, some factors shown to be protective against acid-induced neuronal injury, include lower temperature [63], pharmacological blockade or genetic deletion of ASIC activity [15] or ASIC trafficking [64], i.e. it is established that ASICs play a key role in acidotoxicity. Considering the similar prevalence of ASIC currents, we suggest that the reduced ASIC-mediated current amplitude observed in NMR neurons may be an additional neuroprotective mechanism in NMR brains, alongside the previously described increased hypoxia-inducible transcription factor (HIF1-α) expression [65] or more efficient *in vivo* CO2 buffering [47]. One possible mechanism for the decreased amplitude observed is reduced ASIC plasma membrane trafficking which is known to be modulated by an extracellular acidic environment [64]. However, it is also possible that the reduced acid-induced cell death observed is not ASIC-dependent because although ASIC activation appears to play a major role in neuronal injury [61], several other molecular players are also modulated by a drop in extracellular pH, such as NaVs [27,28] and glutamate receptors [31] and may contribute to the lowered acid-induced cell death observed, for example, here we also show that NMR neurons also have smaller NaV-mediated currents, which may also add a layer of neuroprotection.

## Conclusions

In this work, we describe for the first time the basic electrophysiological properties of NMR neurons in culture and showed that the resting membrane potential of NMR neurons is more hyperpolarized, as well as the amplitude of NaVs being smaller than that of mouse neurons. We then demonstrated that acid-induced currents are present in NMR neurons and are, as in mouse, ASIC-mediated. The key result is that acid-induced cell death is virtually absent in NMR neurons, with reduced ASIC and NaV amplitudes likely contributing to this observation, and thus this is a further adaptation enabling NMR to live in a subterranean, hypercapnic/hypoxic environment.

## Abbreviations

ASIC: acid-sensing ion channel
CNS: central nervous system
DRG: dorsal root ganglion
NaV: voltage-gated Na^+^ channel
NMR: naked mole-rat
TTX: tetrodotoxin

## Declarations

### Ethics

All experiments were conducted in accordance with the United Kingdom Animal (Scientific Procedures) Act 1986 Amendment Regulations 2012 under a Project License (70/7705) granted to E. St. J. S. by the Home Office; the University of Cambridge Animal Welfare Ethical Review Body also approved procedures.

### Consent for publication

N/A

### Availability of data and material

The datasets supporting the conclusions of this article are available in the University of Cambridge Apollo repository: https://doi.org/10.17863/CAM.18704

### Authorsʹ contributions

ZH and ESS designed the study. ZH performed all the experiments and analysed the data. ZH and ESS wrote the manuscript.

## Acknowledgements

Thanks to all members of ESS lab for their useful discussions, and in particular to the assistant staff in the Department of Pharmacology and Animal Facility for their technical help and animal husbandry.

